# Hot, unpredictable weather interacts with land use to restrict the distribution of the Yellow-tailed Black-Cockatoo

**DOI:** 10.1101/2021.04.20.440554

**Authors:** Rahil J. Amin, Jessie C. Buettel, Peter M. Vaughan, Matthew W. Fielding, Barry W. Brook

## Abstract

Conserving nomadic species is challenging due to the difficulty in monitoring their characteristically transient populations, and thereby detecting range-wide declines. An example is the Yellow-tailed Black-Cockatoo (YTBC; *Zanda funerea*), which disperses widely in search of food and is regularly—but sporadically—observed across eastern Australia. Under climate warming, a general southward shift in species distributions is expected in the southern hemisphere, with the extreme southern margins being truncated by an ocean barrier. Given these constraints, we ask whether sufficient refugia will exist for the YTBC in the future, by: (i) modelling habitat relationships within current geographic range of the YTBC based on weather, climate, vegetation, and land use, and (ii) using this framework, coupled with climate-model projections, to forecast 21^st^ century impacts. Intensive land use and high variability in temperature and rainfall seem to most limit YTBC occurrence. In contrast, areas with a cooler, stable climate, and a network of old-growth forests, such as occurs in parts of south-eastern Australia and Tasmania, are most suitable for the species. As Australia becomes progressively hotter under climate change, the preferred bioclimatic envelope of the YTBC is forecast to contract poleward (as a general pattern) and to fragment within the existing range. However, despite an extensive loss of climatically suitable regions, the YTBC might find stable refugia at the southern margins of its geographic range, although continued loss of old-growth forests undermines their nesting potential. Therefore, beyond habitat conservation, creating nesting opportunities within plantation forests would likely be an effective conservation strategy to preserve habitat quality in climate refugia.

## Introduction

Over the last few centuries, anthropogenic conversion of natural habitats and climate change has caused widespread contractions of species distributions globally (Balmford et al. 2002; Brook et al. 2008; Parmesan and Yohe 2003); by 2050 environmental change is predicted to cause at least a 50% loss or shift in the ranges of 400-900 bird species (Jetz et al. 2007). Along with distributional changes across range margins, more frequent extreme-weather events are forecast to lead to die-offs within species’ distributions, especially in hot and arid landscapes such as Australia (McKechnie et al. 2012). Clearly, understanding how birds establish, maintain, and use resources across their ranges is imperative to mitigating further declines.

In Australia, many bird species have evolved nomadic movements in response to an unpredictable and spatiotemporally limiting environment (Rowley 1974; Runge et al. 2015). Resource availability in such landscapes pulsates with rainfall, and opportunistic movements allow birds to capitalise on the transient availability of food, shelter and other resources (Reside et al. 2010). Nomadism is, therefore, strongly entrained to weather conditions, particularly in regions with highly variable inter-annual rainfall patterns (Bateman et al. 2012; Dean and Milton 2001). Granivorous birds are a common functional group in such ecosystems, and many have evolved resource-tracking nomadic behaviour (Elith et al. 2006; Franklin et al. 2000). Indeed, a fifth of Australia’s terrestrial avifauna comprises of granivorous species, but over 30% of these have declined and feature prominently on the list of threatened bird groups (Franklin et al. 2005; Rechetelo et al. 2016).

The conservation of nomadic granivores is spatially challenging because of the difficulty in monitoring their populations, compared to a sedentary species. The extinction vulnerability of vagile species is highly correlated with loss of/shifts in historic geographic ranges (Runge et al. 2015). However, the home ranges of individual nomadic birds are arguably overestimated due to their extensive foraging distances, which can lead to overly pessimistic assessments of the impact of environmental change and subsequently the efficacy of existing management protocols (Runge and Tulloch 2017).

The Yellow-tailed Black-Cockatoo (*Zanda funerea*), hereafter YTBC, is a large, granivorous species, endemic to eastern mainland Australia and Tasmania, including the Bass Strait islands (Higgins 1999; Nelson and Morris 1994). The YTBC is a nomadic parrot which disperses widely in search of food; however, may also become resident during breeding season or under benign conditions (Higgins 1999). The species is widely considered to be ubiquitous throughout this historical range, which likely explains why minimal efforts have been made to investigate the habitat characteristics that limit its distribution or intermittent occurrences within its geographic-range envelope. However, the inadequacy of information on the species’ distribution and habitat requirements means that the prevailing assumption of the ubiquitous status of the YTBC throughout its range might be erroneous, with concomitant implications for conservation management.

Climate change is likely to limit future resource availability across Australia, forcing the YTBC to either adapt or shift to remain in their preferred niche. Poleward shifts in species’ distributions are common biological ‘fingerprints’ of climate change (e.g., Brommer 2004; Conradie et al. 2020; Parmesan et al. 1999). For the YTBC, their southern-range-edge population in south-east Australia and Tasmania is the frontier of the southward redistribution, and perhaps the last refuge, due to a hard ocean boundary. Whether the YTBC can thrive in a future Australian environment depends on the availability of suitable habitat areas (Virkkala et al. 2013), and therefore conservation efforts must be targeted to identify refuge habitats where the YTBC might persist under climate change.

Species distribution models (SDMs; also known as habitat-suitability models) can be used to forecast current and future suitability of habitats based on recorded occupancy patterns, and are particularly useful when species’ physiological responses to environmental change are unknown (Franklin 2010). The correlative approach of SDMs, however, can be limiting when interspecific interactions violate the assumptions of niche constraints, or species are not in approximate equilibrium with their environment due to recent changes in their distribution (Elith et al. 2006; Václavík and Meentemeyer 2012). Despite these limitations, correlative SDMs provide a powerful statistical approach for mapping species distributions and identifying climate refuges – often the first step in conserving nomadic birds (Runge and Tulloch 2017).

Here we model the current and future suitability of habitats for the YTBC across its geographic range in mainland Australia and Tasmania under climate change. We then offer guidelines on how to manage climate-refuge habitats and identify the landscape and vegetation characteristics of most importance in safeguarding the YTBCs’ long-term persistence.

## Methods

### Data collection and processing

We collated occurrence records for the YTBC from the Atlas of Living Australia (download at https://doi.org/10.26197/ala.f3c6b814-e555-486e-8263-964f3686d5bc. Accessed 30 November 2020) and filtered only observations made by eBird and Birdlife Australia (high-quality records) for the decades 1990 to 2020. We removed probable duplicates caused by random YTBC movements by converting point presences into grid presences of 1 km^2^ resolution and retaining only one record per grid cell. Further, potential misidentifications and spatial idiosyncrasies were identified manually and removed. True or confirmed absences were unavailable as they require repeated sampling in space and time. Hence, we generated effort-controlled pseudo-absences by inferring YTBC absence in locations where other bird species have been reported, but not YTBC (i.e., apparent absence for grid cells with observation effort). To achieve this, we reapplied the data collection and processing protocol for occurrence records of all bird species collected between 1990-2020 (Atlas of Living Australia occurrence download at https://bit.ly/3kkwAeE. Accessed 30 November 2020). We then created a buffer zone of a 3 km radius around presence points and retained only those pseudoabsence records outside of the buffer. As undefined spatial extent of pseudo-absences sampling may lead to poor model performance and erroneous predictions (VanDerWal et al. 2009), we also only sampled pseudo-absences within a distance of 50 km from the presence grids – longest distance travelled by a banded YTBC (Higgins 1999). We reduced spatial autocorrelation through data thinning, by re-sampling the presence and pseudo-absence records and removing data points closer than a pre-defined linear distance of *x* (Pearson et al. 2007) (detailed methods in supplementary information). Subsequently, we were left with 4114 presence grids and 4114 pseudo-absence grid cells (i.e., we set a prevalence of 50% for modelling).

### Species distribution modelling

We chose 17 raster layers of environmental predictors that potentially influence distribution of the YTBC (Table 1). We reclassified the vegetation layer to four broad types: rainforest, tall eucalypt forest, woodland, and shrubland/grassland, to avoid overly nuanced classification that would inhibit model fitting. Some species perceive landscape as a continuum of habitat that varies in its relative suitability, and is also modified to different degrees, as compared to conventionally presumed binary selection of habitat or ‘non-habitat’ (Lindenmayer et al. 2003). Therefore, we re-classified the catchment-scale land-use layer to five levels of habitat modification: protected, modified native, plantation, farmland, and human settlement. Due to the lack of available projections on future changes in land use and vegetation-type distribution for Australia, we only forecast climate-induced shifts in the species’ bioclimatic envelope. We used both a high-forcing scenario –one representative of the world’s current CO_2_ emissions trajectory – Representative Concentration Pathway (RCP) 8.5, as well as a mitigation-oriented RCP 4.5 scenario. We used projections from ukmo_hadgem1 general circulation models (GCMs) for both scenarios, projected through to the year 2085 for each weather and climate variable used in the current-day SDM (sourced from BCCVL; bccvl.org.au); these GCMs were selected because of their robust model performance and independence (Evans and Ji 2012). All raster layers were scaled at 1 km resolution to match the occurrence grids. We used a correlative matrix to test for collinearity in current and future environmental layers and removed pairs with high inter-correlation (*r* > 0.7). As modelling algorithms require standardisation of predictor variables, all rasters were centred and normalised using Z-scores prior to analysis.

**Table 1.**
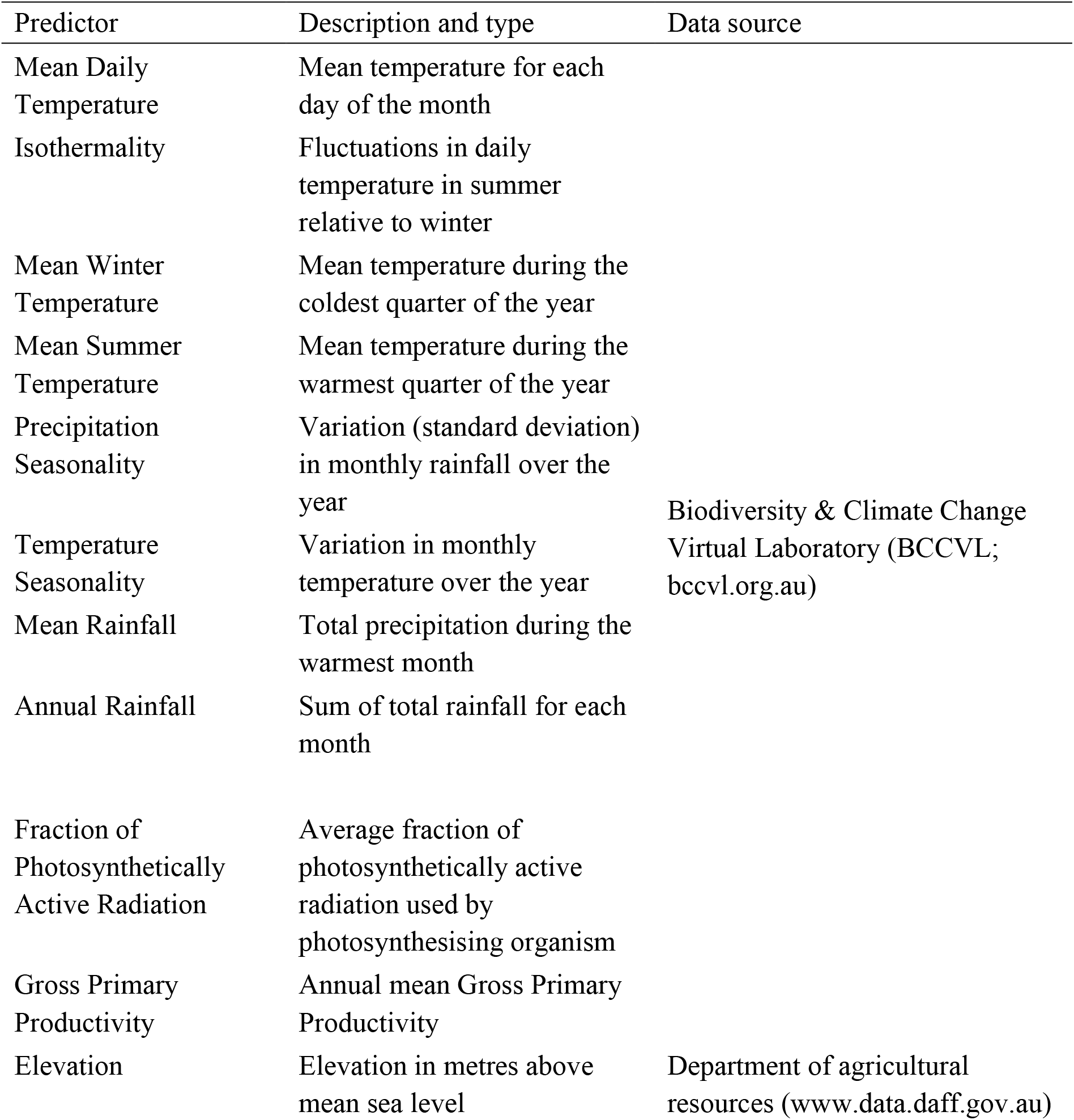
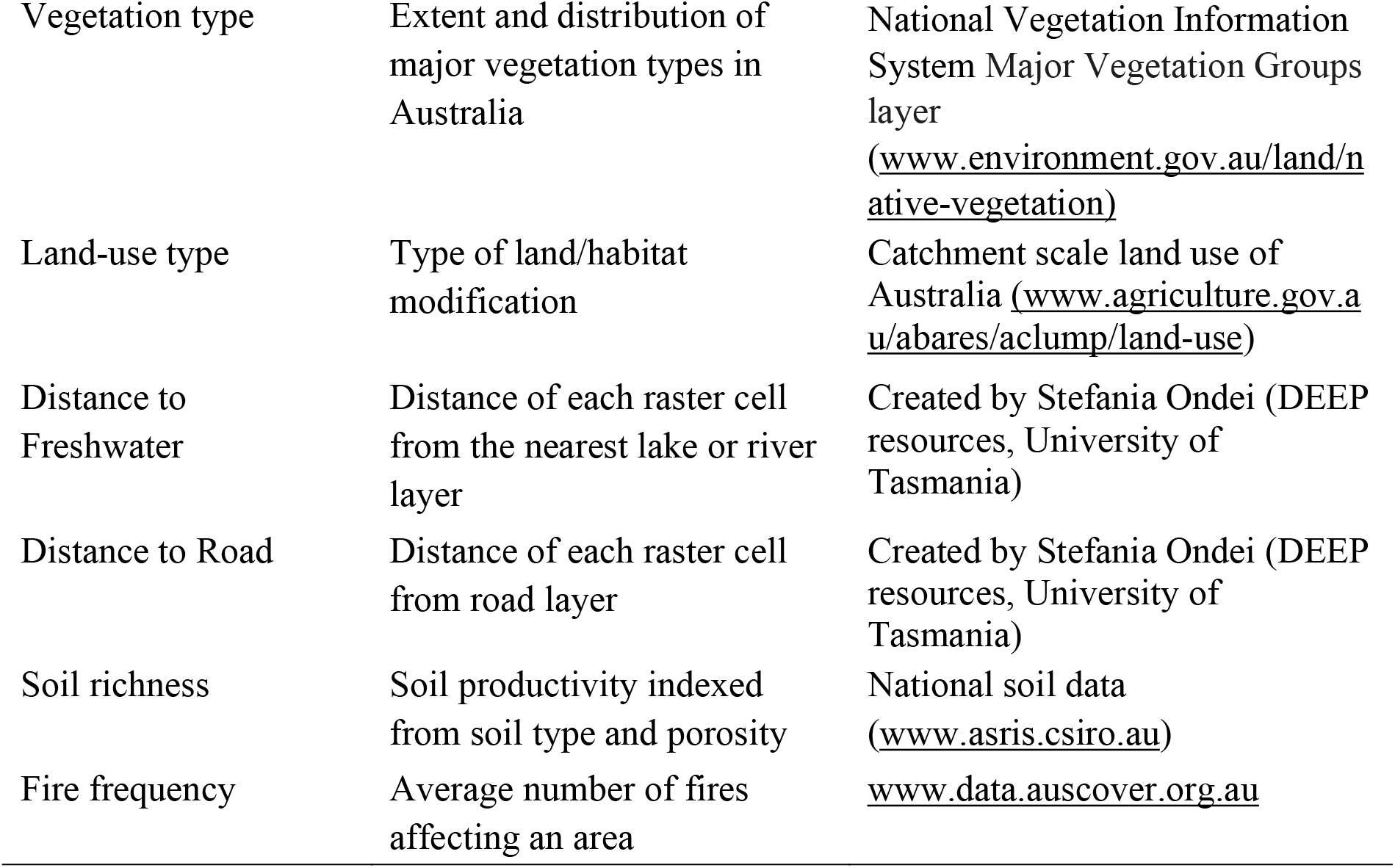
Environmental predictors used to develop species distribution models for the distribution and habitat use of the Yellow-tailed Black-Cockatoo (*Zanda funerea*) in south-eastern Australia.

We assembled the SDM in the R package `sdm` (Naimi et al. 2016) using an ensemble of four machine-learning-model algorithms: generalised linear model (GLM), generalised additive model (GAM), random forest classification (RF), and gradient boosted machine (GBM); the former two are additive regression-type approaches, the latter two are decision trees. We then used this ensemble to forecast the current habitat suitability map and future changes in bioclimatic envelope for the YTBC in Australia. We chose the Area Under the Curve (AUC) of Receiver Operating Characteristic (ROC) curve as a metric to assess the accuracy of SDMs due to its advantage in obtaining a threshold independent measure for evaluating both false-positives and false-negatives (Allouche et al. 2006). We used repeated k-fold cross validation (with *k* = 25, and ten repetitions) in the R package `caret` to evaluate model performance and accuracy and to tune the model parameters until, with the target of a maximum AUC value for the test (k-fold hold-out) data. AUC values were also used for model refinement (variable selection/rejection), where only predictor variables that provided the most accurate and parsimonious predictions were retained. We then passed the tuned parameters to package `sdm` for ensemble predictions with weighting by test-score AUC (Allouche et al. 2006; Marmion et al. 2009).

## Results

### Habitat use by the YTBC

Of the 17 predictors, vegetation type, land-use type, productivity (i.e., fraction of Photosynthetically Active Radiation absorbed (fPAR), seasonality in temperature, mean daily temperature, mean temperature in summer, and seasonality in rainfall provided the most parsimonious and accurate model projections of YTBC occurrence (ensemble AUC = 0.88; where a random expectation = 0.5 and a perfect discriminator = 1). Seasonality in inter-annual weather patterns was the strongest predictor of the probability of YTBC occurrence (Figure 1), implying that the species prefers stable weather conditions; high variability in both temperature and rainfall caused a sharp drop in the log-likelihood of species’ presence (Figure 1). The YTBC also preferred cooler areas where the mean daily temperature and summer temperatures were relatively lower (Figure 1). Consequently, the species was predicted to preferentially use rainforests (base odds ratio (OR) = 0.95) over other vegetation types (Table 2), with grassland and shrubland occupancy being lowest (OR = −1.25 relative to the base; Table 2). In terms of land use, the YTBC was most likely to occur in National parks and forestry areas (base OR = 0.95), while also being common in urban areas (OR = 0.86; Table 2). Their apparent ubiquity in urban areas, however, is likely due to observer bias. Interestingly, we found a positive correlation between occurrence probability and pine plantations (OR = 0.58; Table 2). The clearance of the most suitable YTBC habitat areas for agriculture (OR = −0.72; Table 2), grazing, and intensive silviculture (OR = −0.88; Table 2) acts to limit the likelihood of occurrence, constraining their distribution within Australia (Table 2; Figure 2). Finally, YTBC was predicted to prefer low-productivity habitats (Figure 1).

**Table 2.** Generalised additive model (GAM) response curves for the change in occurrence probability of the Yellow-tailed Black-Cockatoo (*Zanda funerea*) with habitat modification and vegetation type (SE = standard error, Std. coef = standardised regression coefficients).

| Variable | Odds ratio | SE | Std. coef. |
| --- | --- | --- | --- |
| National Parks and Reserves + Rainforest | 0.95 | 0.16 | 0 (base) |
| Modified native | -0.88 | 0.06 | -0.28 |
| Farmland | -0.72 | 0.07 | -0.20 |
| Plantation | 0.58 | 0.11 | 0.10 |
| Human settlement | 0.86 | 0.19 | 0.09 |
| Wet forest | 0.41 | 0.17 | 0.05 |
| Woodland | -1.21 | 0.17 | -0.14 |
| Grasslands/shrubland | -1.25 | 0.17 | -0.14 |

**Figure 1.**
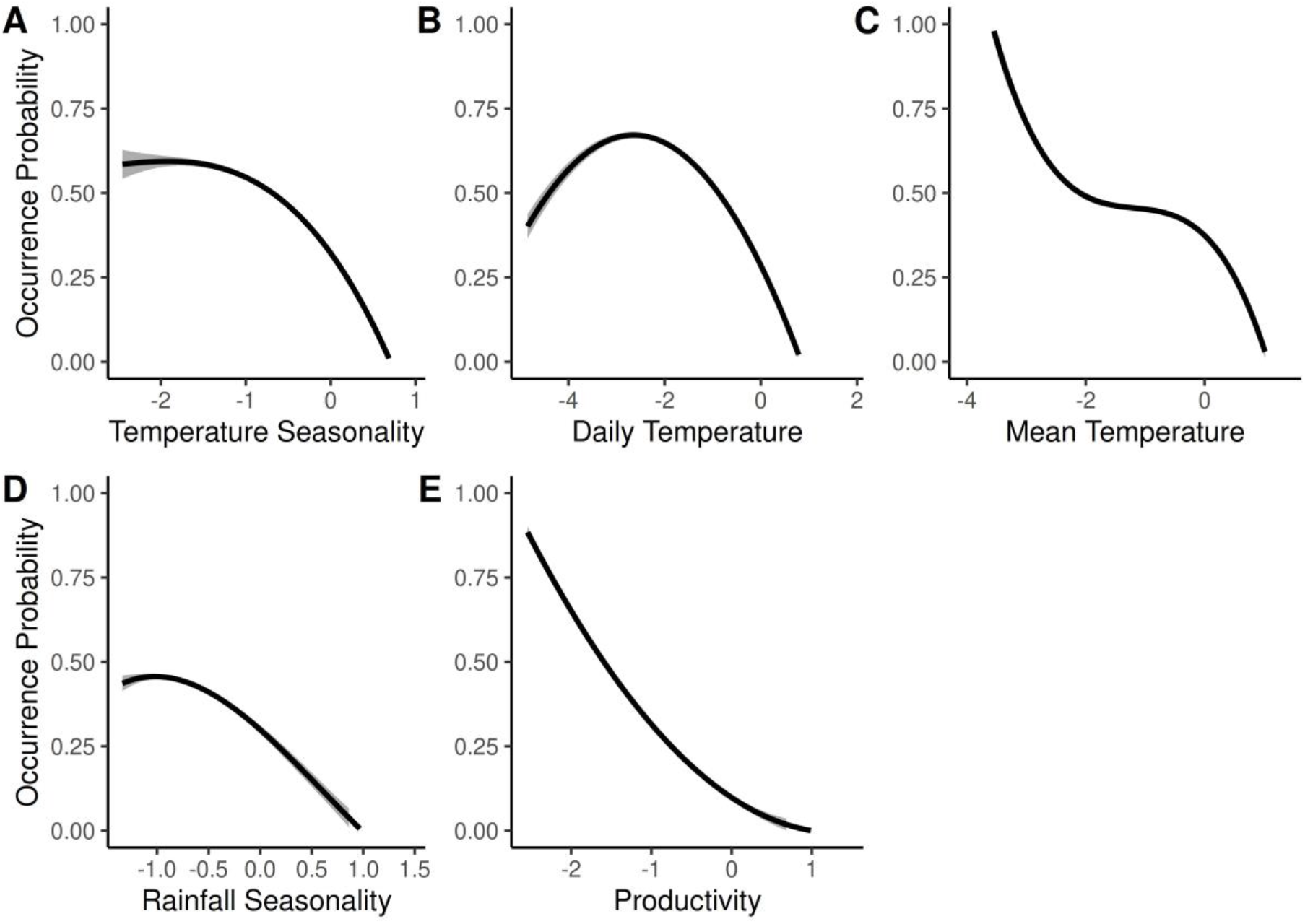
Generalised additive model response curves for change in the occurrence probability of the Yellow-tailed Black-Cockatoo (*Zanda funerea*) with climate and weather conditions [seasonality (standard deviation) in temperature (A), mean daily temperature (B), mean summer temperature (C), seasonality in precipitation (D)], and productivity (i.e., the Fraction of Photosynthetically Active Radiation (E).

**Figure 2.**
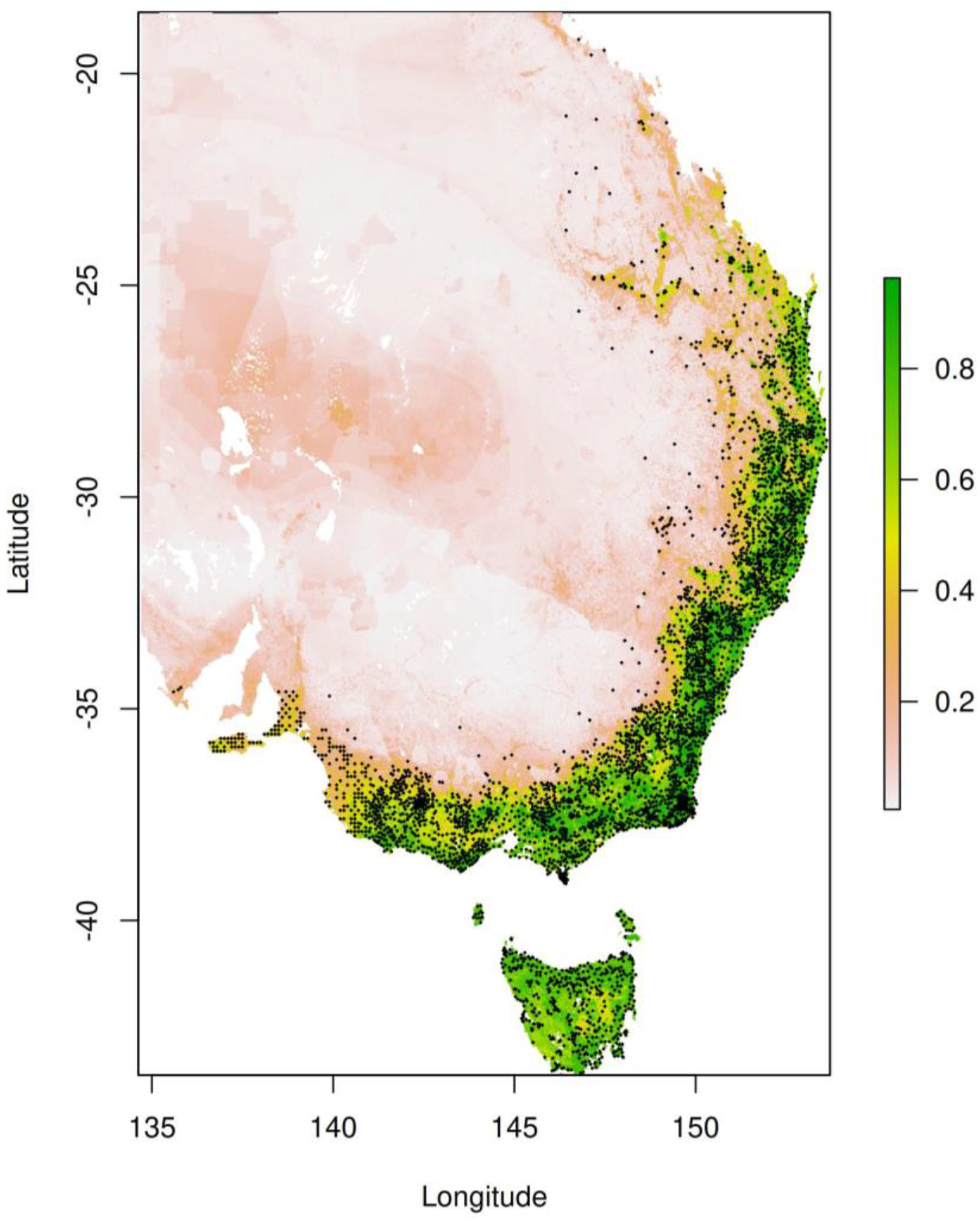
Species distribution model ensemble projections (from statistical into geographic space) for current suitable habitats for the Yellow-tailed Black-Cockatoo (*Zanda funerea*). The colour scale represents the relative habitat suitability index, scaled from 0 to 1 (i.e., from the least- to most-suitable grid cell), where the green areas represent highly suitable habitats and white areas are unsuitable. The black dots are historical occurrences of the YTBC.

### Current suitable habitats and climate induced shifts

The eastern seaboard of mainland Australia, Tasmania, and the Bass Strait Islands were projected to encompass the current suitable habitats for the YTBC—congruent with the geographic distribution of largely intact temperate forests (Figure 2). In contrast, most of the inland regions of mainland Australia were unsuitable due to the high variability, hot weather and the absence of preferred vegetation (Figure 1 and 2; Table 2) (Jackson 1999). Regardless of the emission scenario, climate change is forecast to cause poleward and eastwards contractions in, and fragmentation of, the species’ climatic niche by the year 2085, albeit to a varying degree in different regions (Figure 3). Furthermore, these losses will be more evident in highly suitable areas in southern Queensland and eastern New South Wales (Figure 3 and 4). Despite the extensive losses of their climatic niche with western inland regions, the YTBC should still find preferred conditions at the eastern mainland edge (especially higher-elevation zones) and the southern margins of its geographic range (Figure 2 and 4).

**Figure 3.**
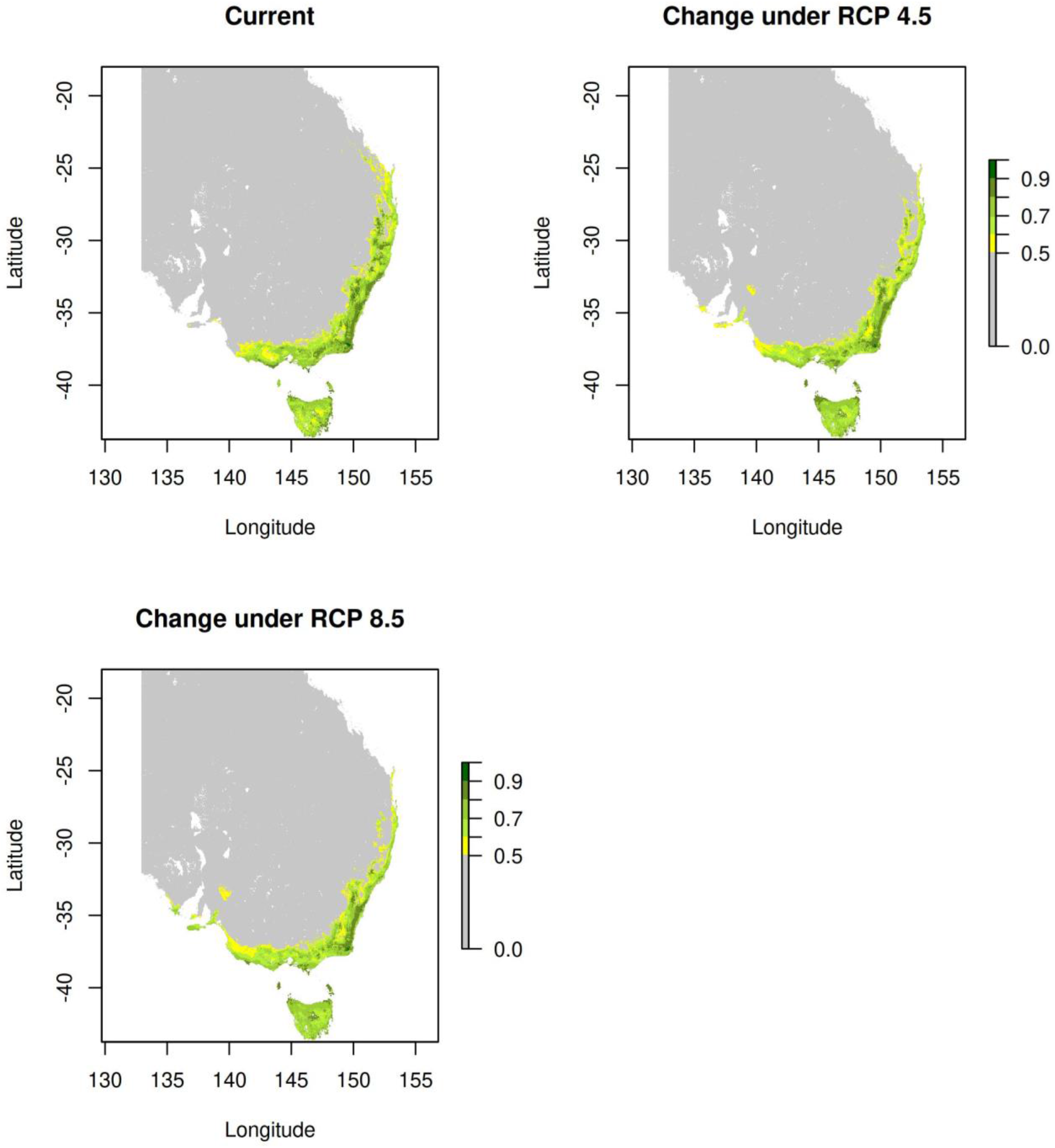
Forecast change in the suitability of Yellow-tailed Black-Cockatoo’s (*Zanda funerea*) climatic niche across Australia from current distribution to the year 2085 under the worst-case (RCP 8.5) and moderate (RCP 4.5) climate-change scenario considered by global climate models. The colour scale represents the relative climatic suitability index, scaled from 0 to 1 (i.e., from the least- to most-suitable grid cell), where the dark green areas represent highly suitable habitats (HSI > 9), yellow areas represent sub-optimal areas (HSI > 0.5), and grey areas are unsuitable (HSI < 0.5)

**Figure 4.**
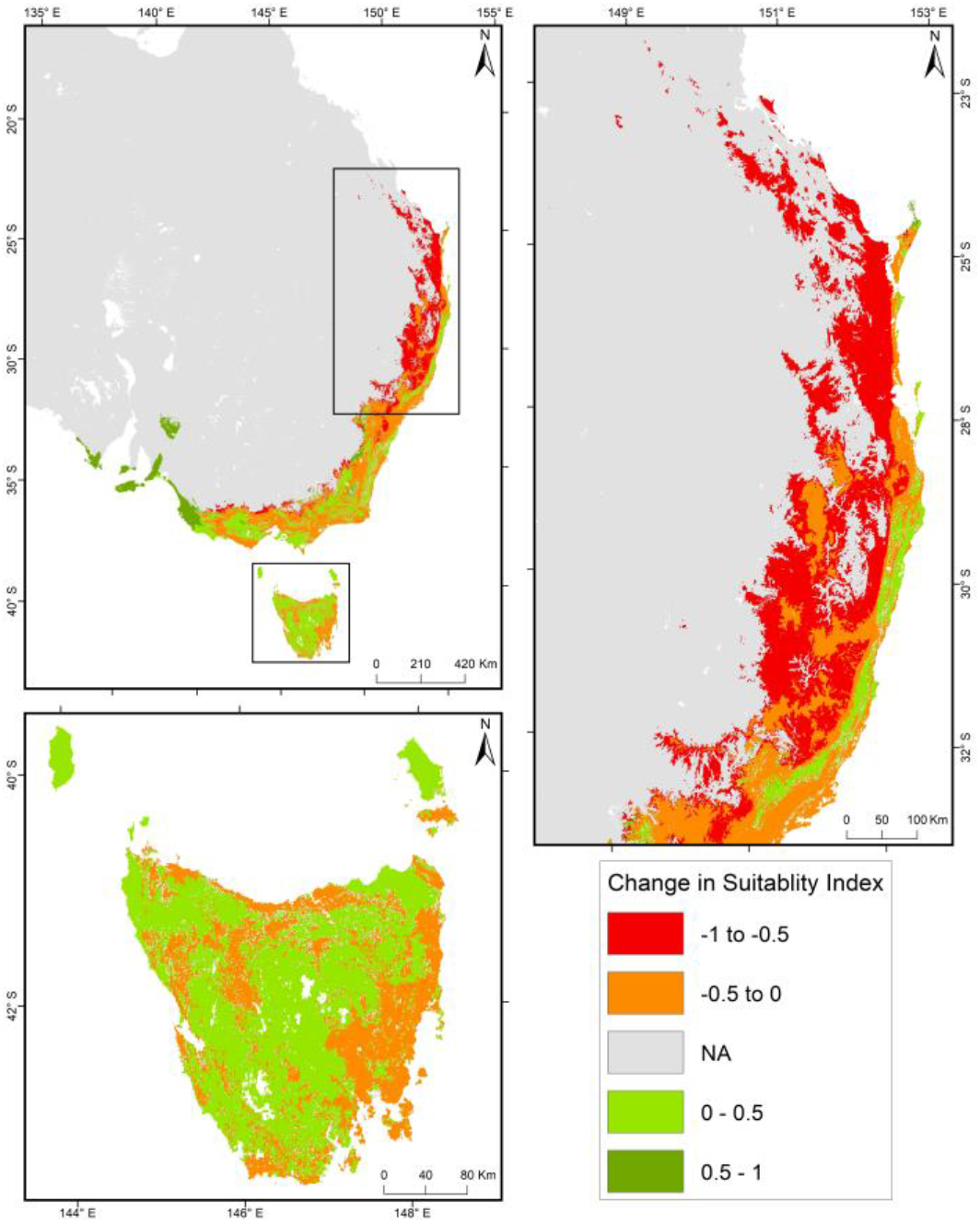
Loss or gain in climatically suitable habitats (i.e., areas with model-predicted Habitat Suitability Index HSI > 0.5) for the Yellow-tailed Black-Cockatoo (*Zanda funerea*) by the year 2085 under the worst-case climate-change scenario (RCP 8.5). Insets show two areas of particular importance— southern Queensland and eastern New South Wales in the north of the range, and Tasmania in the south. The colour scale represents the change in suitable climatic suitability index, scaled from −1 to +1, where the dark red areas represent areas with prominent loss in climatic suitability (HSI _(2085)_ – HSI _(2020)_ < - 0.5) and dark green represents the areas with increase in climatic suitability (HSI _(2085)_ – HSI _(2020)_ > 0.5).

## Discussion

Hot climate, unpredictable weather and intensive habitat conversion are the factors most limiting the current distribution of YTBC. As a result, the YTBC is most prevalent across eastern seaboard of mainland Australia, parts of Tasmania, and Bass Strait Islands due to these regions supporting areas with cooler, more stable weather patterns, and an extensive network of rainforests within national parks and reserves. However, historically, the species has also ranged over the region to the west of the Great Dividing Range.

The YTBC has evolved in an environment characterised by an ephemeral abundance of resources that boom and bust with weather patterns (Dean and Milton 2001; Franklin et al. 2000; Rechetelo et al. 2016). Extreme events, such as prolonged heat waves coupled with rainfall deficits, may be more common in highly variable weather and may indirectly limit resource availability (Reside et al. 2010). Furthermore, fluctuating temperatures can also disrupt the physiological functions of individuals (Butler and Woakes 1990; Chaplin 1976), affecting their resource-acquisition ability. Consequently, stable weather provides relatively secure and predictable habitats for the YTBC.

The flocks that engage in frequent, energetically expensive foraging movements, risk mortality and must locate resources before their stored energy drops below critical levels (Bennetts and Kitchens 2000; Pyke et al. 1977). Consequently, the proximity of habitat directly influences the energy intake-to-expenditure ratio of the foraging individuals (Pyke 1984). Increasing variability in climate—leading to frequent extreme weather events from climate change—has the general effect of increasing the distance between suitable patches and thereby degrading the overall suitability of the landscape (Reside et al. 2010). A degraded landscape might force YTBC flocks to increase foraging movements to meet their minimum energy demands, with concomitant fitness penalties.

The low moisture content of seeds (their primary food source) forces YTBC to rely on close proximity to perpetual water sources to satisfy their metabolic water requirements, explaining their preference for cooler-weather conditions and rainforests. Increased air temperatures are predicted to cause large increases in thermoregulatory water requirements, particularly in granivores (McKechnie and Wolf 2010). In the last two decades, dehydration during heat waves or extremely hot days (> 45°C) has caused mass fatalities in other nomadic granivores, including Budgerigars (*Melopsittacus undulatus*), Zebra Finches (*Taeniopygia guttata*), and the endangered Carnaby’s Black-Cockatoo (*Zanda latirostris*) (McKechnie et al. 2012). The YTBC is also vulnerable to dehydration and high energy expenditure under a warmer climate, due to increasingly frequent foraging movements that might not be compensated by resource uptake in a simultaneously degraded landscape.

The YTBC is a large, long-lived bird with a low rate of reproduction, and relies on hollows of old-growth eucalypt trees for nesting (Nelson and Morris 1994). Tree hollows are formed by slow processes and take many years to reach YTBC preferred dimensions (Saunders et al. 1982). The national parks and reserves in species’ geographic range encompass extensive networks of rainforests with old-growth eucalypts that are used preferentially by the YTBC (Mendel and Kirkpatrick 2002; Nelson and Morris 1994). Therefore, the continued historical clearance of old-growth eucalypt- and rain-forests in YTBC-suitable habitats for use by agriculture and short-rotational silviculture will reduce the number of hollow-bearing trees and nesting opportunities for the YTBC (Mendel and Kirkpatrick 2002). When paired with die-offs from predicted increases in the frequency and severity of heat waves, along with the increased vulnerability of hollow trees to fire-related mortality, the YTBC is in danger of declines due to supressed breeding success and rare but damaging mortality events.

At present, the east-coast elevated areas and southern edge of mainland Australia and Tasmania provide the YTBC a geographical respite from these threats. The prevalence potential foraging and nesting competitors of the YTBC, being other Cacatuidae species and nest predators like Brown Tree Snakes (*Boiga irregularis*), in mainland Australia may limit the nesting and foraging potential of habitats in Victoria and New South Wales (Higgins 1999; Wiles et al. 2003). These pressures, however, are likely to remain low in Tasmania, providing YTBC an ideal refuge under climate change (Nascimento et al. 2011). Indeed, as Bass Strait will inhibit fluid niche-tracking movements of the mainland population, in the future Tasmania might be the main (or only) sanctuary for the YTBC.

We have identified potentially suitable refuge regions, which are, or in the future are likely to become, climatically suitable for the YTBC. However, the quality of these climate refuge habitats will vary locally, and not all identified habitats may meet the YTBCs’ resource requirements (e.g., cleared forests or farmland). Therefore, despite south-east Australia and Tasmania being strongholds climatically, degraded nesting and foraging habitats will impede the species’ ability to adapt and seek refuge. It is therefore crucial that conservation actions are targeted to protect suitable climate-refuge habitats from further loss.

Regarding this last point, the efficacy of YTBC conservation actions ultimately depends on the maintenance of a self-sustaining population. Therefore, managers must aim to conserve potential nesting sites in old-growth native forests and retain mature trees in logging coupes. Further, the nesting potential of anthropogenic vegetation types that are used extensively by YTBC for feeding, such as pine plantations, can be enhanced using artificial nesting hollows that meet the YTBC preferred dimensions. While we found only a weak positive correlation, YTBC has been frequently observed foraging on *Pinus radiata* seeds (Higgins 1999). Therefore, pine plantations with artificial nesting hollows could emulate native habitats suitable for both foraging and nesting, extending the total habitat area available to the YTBC. The further provision of water might reduce the risk of dehydration in foraging individuals. The maintenance of a viable, self-sustaining population of YTBC is most probable through the spatial orchestration of conservation of existing suitable habitat areas and the provision of nesting hollows and foraging resources in non-native vegetation and land-use types that are otherwise climatically suitable.

The stochasticity in movement patterns and distributional dynamics of nomadic YTBC, leading to apparent ubiquitous occurrences, make it difficult to estimate their true extinction vulnerability. While we have met this challenge in part—with debiasing, thinning and careful selection of absence data—our predictions are nevertheless based on observations pooled across time, and therefore represent averaged habitat use by the YTBC. Consequently, the predicted decline in habitat suitability might be underestimated. Furthermore, their predicted prevalence in areas of high human density is likely to have resulted from some residual spatial autocorrelation, despite our best efforts to mitigate this observational bias. Although citizen-science data (such as birdwatching records) provide essential information on the occurrence of an understudied species, SDMs based on these data must be applied and interpreted with caution. Ultimately, accounting for data biases alongside the sporadic nomadic movements of flocks and considering the likelihood of success of management interventions across large landscapes, will be most useful in informing conservation solutions for the YTBC that are effective, cost-efficient, and robust to uncertainty under global change.

## Acknowledgements

Thank you, all the authors, for their contributions to this work. The authors thank the editor, associate editor and reviewers for their suggestions and comments, which has resulted in a much-improved paper.

## Supplementary information

### Debiasing spatial autocorrelation

Spatial autocorrelation or spatial bias is a common problem in citizen-science data on species occurrences, as these data are collected preferentially from easily accessible or heavily visited areas (e.g., roads or settlement areas). As such, they are not a random and representative sample of a species’ distribution (Barve et al. 2011). Non-random sampling can lead to environmental bias, where higher sampling effort in some regions results in overrepresentation of environmental conditions in those regions. Therefore, data debiasing is required to maximise the signal-to-noise ratio when fitting a species distribution model (SDM) to citizen-science data.

Spatial thinning of data provides one solution, wherein the occurrence records are filtered in regions with high data density/clustering while retaining data where density is low. Subsequently, this process mitigates overrepresentation of certain localities by thinning observation clusters. Therefore, we reduced spatial bias by re-sampling the occurrence (presence and pseudo-absence) grids used for building our models, and removing occurrence records closer than a pre-defined (minimum linear) nearest minimum-neighbour distance (NMD), *x* (Pearson et al. 2007). The optimal NMD (*x*) varies with species and study regions, and is typically user-defined due to a lack of available systematic approaches (Aiello-Lammens et al. 2015). Occurrence records from regions of low human density are least likely to be spatial autocorrelated. Therefore, the average number of records per low-human-density grid (see below) provides the lower threshold number of records that spatially autocorrelated areas (with higher human density) should have recorded. Consequently, optimal NMD must be adjusted (tuned) such that the threshold average can be reached when thinning records in spatially autocorrelated areas.

We achieved this by stratifying Australia into low-, medium-, and high-density 25km^2^ grid cells. To ensure spatially uncorrelated data in low-density grids, we removed records closer than *x* = 4 (retains most records while reducing clustering). Further, we calculated the threshold number of re-samples per grid (*h*) using:

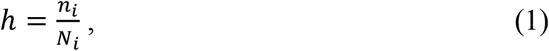

where *n_i_* represents the number of samples in low-density grids and *N_i_* is the number of low-density grids (these areas, having the fewest observations per unit area, sets the lower bound for thinning of the observations). Subsequently, we used the threshold value (*h*) to estimate the maximum number of re-samples to be taken from medium and high-density grids, using the formula:

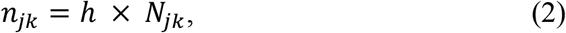

where *n* is the maximum number of re-samples from medium *j* or high *k* density grids, and *N* is the number of medium *j* or high *k* density grids. We implemented the NMD approach using the ‘thin’ function in r package `spThin` (Aiello-Lammens et al. 2015) and randomly re-sample the YTBC occurrence (presence/pseudo-absence) records. We repeated this 12 times per sampling run and adjusted the *x* for high- (*x*_k_ = 12) and medium-density grids (*x*_j_ = 8) to achieve the maximum number of re-samples (*n*_jk_). The dataset with the most records was retained from each grid type (i.e., high, medium, and low) and collated into the final dataset.

### Pseudo-code for debiasing spatial autocorrelation

1. Calculate mean human density for each 25 km^2^ grid cell.
2. Plot data distribution of mean human density and estimate quartiles.
3. Split grids by mean human density as: high (mean density > 0.75 quartile), medium (0.25 < mean density < 0.75), and low (mean density < 0.25 quartile).
4. Split species data by grids into high-, medium-, low-density occurrence records.
5. Thin low-density records by *x* = 4.
6. Calculate threshold values (*h*) = number of low-density occurrence records (*n_i_*) / number of low-density grids (*N_i_*).
7. Calculate maximum number of resamples from high-density grids (*n_k_*) = threshold value (*h*) × number of high-density grids (*N_k_*); and maximum number of resamples from medium-density grids (*n_j_*) = threshold value (*h*) × number of medium-density grids (*N_j_*).
8. Thin high density occurrence records using distance *x*_k_ = 12 and 12 repetitions to reach maximum number of desired records (*n_k_*). Select the dataset with the highest number of resamples.
9. Thin medium-density occurrence records using distance *x*_j_ = 8 and 12 repetitions to reach maximum number of desired records (*n_j_*). Select the dataset with the highest number of resamples.
10. Collate high-, medium-, and low-density occurrence dataset into final species data.

### Species response curve

Generalised additive models (GAMs) are appropriate for modelling complex ecological responses (Elith et al. 2006), using non-parametric, data-defined smoothing splines to fit non-linear relationships. Therefore, we implemented GAMs using binomial error distribution with a logit link in the R package `caret` to predict the probability of habitat use by the YTBC.

**Table (SI).** Out-of-sample predictions for species distribution models fitted to the presence-pseudo-absence occurrence data of the Yellow-tailed Black-Cockatoo (*Zanda funerea*). For each model-species combination, values for Area Under the Curve (AUC) for Receiver Operating Characteristic (ROC) and Test Skill Statistic (TSS) are shown. GLM = generalised linear models; GAM = generalised additive model; RF = random forests; GBM = gradient boosted machine.

|  | GLM | GAM | RF | GBM | ensemble |
| --- | --- | --- | --- | --- | --- |
| AUC | 0.85 | 0.87 | 0.88 | 0.88 | 0.87 |
| TSS | 0.54 | 0.57 | 0.61 | 0.60 | 0.60 |

